# Evolution of the jasmonate ligands and their biosynthetic pathways

**DOI:** 10.1101/2023.02.03.526968

**Authors:** Andrea Chini, Isabel Monte, Angel M. Zamarreño, José M. García-Mina, Roberto Solano

**Affiliations:** Plant Molecular Genetics Department, Centro Nacional de Biotecnologia-CSIC (CNB-CSIC), 28049 Madrid, Spain; Department of Environmental Biology, Bioma Institute, University of Navarra, Navarra 31008, Spain; Center for Plant Molecular Biology (ZMBP), University of Tübingen, 72076 Tübingen, Germany

## Abstract

Jasmonates are phytohormones that regulate multiple aspects of plant development and responses to stress, activating a conserved signaling pathway in land plants. The characterization of jasmonates biosynthetic and signaling pathways revealed that (+)-*7-iso*-JA-Ile (JA-Ile) is the ligand for the COI1/JAZ receptor in angiosperms, where jasmonates are synthesized through the OPR3-dependent or OPR3-independent pathways. More recently, studies on different model species identified dn-*cis*-OPDA, dn-*iso*-OPDA and Δ^4^-dn-*iso*-OPDA as the ligands of the COI1/JAZ receptor in the liverwort *Marchantia polymorpha*, and a receptor-independent role for several jasmonates in streptophytes. To understand the distribution of bioactive jasmonates in the green lineage and how their biosynthetic pathways evolved, we combined phylogenetic analyses and jasmonates metabolomics in representative species from different lineages. We found that both OPDA and dn-*cis*-OPDA are ubiquitous in land plants and present also in charophyte algae, underscoring their importance as ancestral signalling molecules. In contrast, JA-Ile biosynthesis emerged within lycophytes coincident with the evolutionary appearance of JAR1 function. We show that JA biosynthesis mediated by OPR1/OPR2 appeared in charophytes most likely as a degradation pathway of OPDA/dn-*cis*-OPDA before OPR3 emergence. Therefore, our results demonstrate that the OPR3-independent JA biosynthesis pathway is ancient and predates the evolutionary appearance of the OPR3-dependent pathway. Moreover, we identified a negative correlation between dn-*iso*-OPDA and JA-Ile in land plants which supports that dn-*iso*-OPDA is the relevant form of the hormone perceived by COI1/JAZ in bryophytes and lycophytes.

## Introduction

Jasmonates are lipid-derived hormones that play essential roles in plant development and defense against herbivores and necrotrophic pathogens (Howe *et al*., 2018; Wasternack & Feussner, 2018). Extensive studies on angiosperms, mainly *Arabidopsis thaliana*, identified (+)-7-*iso*-JA-Ile as the ligand of the jasmonate co-receptor complex formed by the F-box coronatine insensitive 1 (COI1) and the jasmonate ZIM domain (JAZ) repressors (Xie *et al*., 1998; Chini *et al*., 2007; Thines *et al*., 2007; Fonseca *et al*., 2009; Sheard *et al*., 2010). More recently, studies in the bryophyte *Marchantia polymorpha* and the lycophyte *Huperzia selago* identified two isomers of dn-OPDA: dn-*cis*-OPDA and dn-*iso*-OPDA as the COI1/JAZ ligands in both species, and Δ^4^-dn-*iso*-OPDA as an additional ligand in *M. polymorpha* (Monte et al., 2018; Kneeshaw et al., 2022; Monte et al., 2022). However, whether these dn-OPDA-related molecules (collectively named here as dn-OPDAs) are also present and may act as bioactive jasmonates in other plants species has not been addressed yet. Even though JA-Ile and dn-OPDAs are perceived by orthologous COI1/JAZ co-receptor complexes, differences in ligand binding were attributed to a single amino acid change in COI1 that alters the binding pocket size between bryophytes and vascular plants (Bowman et al., 2017; Monte et al., 2018). More recently, we discovered that specific residues within the JAZ degron contribute to ligand specificity together with COI1 (Monte *et al*., 2022).

Hormone perception leads to the degradation of the JAZ repressors and activation of conserved MYC transcription factors in land plants (Chini et al., 2007, 2009; Thines et al., 2007; Sheard et al., 2010; Fernández-Calvo et al., 2011; Monte et al., 2018, 2019; Peñuelas et al., 2019). Jasmonates induce similar transcriptional reprogramming comprising genes related to wounding and secondary metabolism in both *M. polymorpha* and *A. thaliana* (Chini et al., 2016; Howe et al., 2018; Monte et al., 2018, 2019; Peñuelas et al., 2019). However, while JA-Ile regulates defense, growth, and fertility, dn-OPDA regulates defense and growth but not fertility in *M. polymorpha* (Monte et al., 2018, 2019; Peñuelas et al., 2019). In addition, OPDA and dn-*cis*-OPDA exhibit signaling properties independently on COI1, regulating thermotolerance due to their reactive electrophilic activity in streptophytes such as the charophyte alga *Klebsormidium nitens*, the liverwort *M. polymorpha* or the eudicot *Arabidopsis thaliana* (Monte *et al*., 2020). Interestingly, OPDA, dn-OPDAs and JA-Ile share the initial biosynthetic steps of the octadecanoid/hexadecanoid pathway up to the formation of OPDA/dn-*cis*-OPDA, which are indeed precursors of JA-Ile in angiosperms. Briefly, the fatty acids linolenic acid (18:3), hexadecatrienoic acid (16:3) and eicosapentanoic acid (20:4) are released from the chloroplast membranes by phospholipases and processed into OPDA, dn-*cis*-OPDA or Δ^4^-dn-OPDA, respectively, by the coordinated action of lipooxygenases, allene oxide synthase (AOS) and allene oxide cyclase (AOC) (Weber et al., 1997; Wasternack & Song, 2017; Kneeshaw et al., 2022). In angiosperms OPDA and dn-*cis*-OPDA are then transported to the peroxisome where one key enzyme, OPDA-reductase 3 (OPR3) reduces the double bond in the cyclopentenone ring of both OPDA and dn-*cis*-OPDA, producing OPC-8 and OPC-6, respectively (Schaller *et al*., 2000; Wasternack & Feussner, 2018). These molecules undergo three or two rounds of beta-oxidation to give rise to JA (Li *et al*., 2005). Synthesis of JA-Ile requires conjugation of JA to Ile, which occurs in the cytosol and is catalysed by Jasmonate Resistant 1 (JAR1) (Staswick & Tiryaki, 2004). Minor contribution of other enzymes from the same Glycoside Hydrolase (GH3) family and conjugation of JA to other amino acids have been reported (Staswick & Tiryaki, 2004; Delfin et al., 2022). JA-Ile or dn-OPDAs are translocated into the nucleus and perceived by COI1/JAZ in *A. thaliana* or *M. polymorpha/H.selago*, respectively. Besides the canonical jasmonate biosynthetic pathway utilizing the peroxisomal OPR3, there is an alternative biosynthetic pathway dependent on the cytosolic OPR3 homologs OPR1 and OPR2 (Strassner *et al*., 2002; Chini *et al*., 2018). While OPR3 accepts OPDA as a substrate due to two key residues (F74 and H265), OPR1/OPR2 lack these two amino acids and hence cannot reduce OPDA directly (Breithaupt *et al*., 2009). Instead of being reduced by OPR3, OPDA can alternatively enter subsequent beta-oxidation cycles and give rise to dn-OPDA, tetranor-OPDA (tn-OPDA) and 4,5-didehydro-JA (4,5-ddh-JA), which is a substrate for OPR1 and OPR2 (Chini et al., 2018; Monte et al., 2018). Thus, OPR1 and OPR2 can catalyze the reduction of 4,5-ddh-JA to JA in the cytosol independently on OPR3 (Chini *et al*., 2018). OPR3 has been proposed to emerge in vascular plants, whereas OPR1/OPR2 origin is more ancient (Bandara et al., 2012; Pratiwi et al., 2017; Chini et al., 2018; Monte et al., 2018). Although the OPR3-dependent pathway is the main pathway for JA-Ile synthesis in *A. thaliana*, the evolutionary origin, the relative importance of each pathway and how broadly are OPR3-dependent and -independent pathways represented along the phylogeny of plants has not been addressed yet.

In addition to *A. thaliana*, the JA-Ile precursors OPDA and dn-*cis*-OPDA have been described in different non-vascular species including the bryophytes *Physcomitrium patens* (formerly *Physcomitrella patens)* and *M. polymorpha* and the charophyte alga *K. nitens*, suggesting that the origin of OPDA/dn-OPDA biosynthesis predates land plant colonization (Weber et al., 1997; Stumpe et al., 2010; Koeduka et al., 2015; Chini et al., 2018; Monte et al., 2018, 2020). Conversely, the presence of JA/JA-Ile in algae, bryophytes and lycophytes is controversial (Scholz et al., 2012; Beilby et al., 2015; Koeduka et al., 2015; Záveská Drábková et al., 2015; Monte et al., 2018; Nishiyama et al., 2018; Široká *et al*., 2022; Resemann *et al*., 2023) and it has been proposed that JA-Ile biosynthesis emerged in lycophytes (Pratiwi et al., 2017). To clarify the evolution of the JA-Ile biosynthetic pathway and the distribution of bioactive jasmonates in the green lineage, we performed phylogenetic analyses and oxylipin metabolomic profiling in representative species covering the main clades of green algae and land plants under control or stress conditions. We confirmed that OPDA and dn-*cis-*OPDA emerged in charophyte algae and are ubiquitous in land plants. We showed that JA biosynthesis mediated by OPR1/OPR2 appeared in charophytes most likely as a degradation pathway of OPDA/dn-*cis*-OPDA before OPR3 emergence. Therefore, our results demonstrate that the OPR3-independent JA biosynthesis pathway is ancient and predates the evolutionary appearance of the OPR3-dependent pathway. We confirmed that the biosynthesis of JA-Ile emerged within lycophytes although its accumulation kinetics does not seem compatible with a role as stress hormone in this lineage. Remarkably, we detected a negative co-relation between dn-*iso*-OPDA and JA-Ile in all land plant lineages except in *S. moellendorffii*. Our results support that the function of dn-*iso*-OPDA as a COI1/JAZ ligand is not restricted to *M. polymorpha*, and that in bryophytes and lycophytes dn-*iso*-OPDA represents the analogous hormone to JA-Ile in other vascular plants.

## Results

### OPDA and dn-*cis*-OPDA emerged in charophytes and are ubiquitous in land plants

To investigate the evolution of jasmonates biosynthesis and the identification of relevant molecules mediating jasmonates responses to stress we performed oxylipin metabolomic analyses on representative species within the green lineage after heat shock or wounding. In addition to previously analyzed model organisms (the charophyte *K. nitens*, the liverwort *M. polymorpha* and the eudicot *A. thaliana*), we selected one chlorophyte *(Chlamydomonas reinhardtii*), one hornwort (*Anthoceros agrestis*), three mosses (*P. patens, Polytrichastrum formosum, Dicranum scoparium*), four lycophytes (*Selaginella moellendorffii, H. selago, Lycopodium clavatum, Diphasiastrum alpinum*), one horsetail (*Equisetum hyemale*), one fern (*Nephrolepis exaltata*) and one monocot (*Triticum durum*). We observed that the chlorophyte *C. reinhardtii* was devoid of all the oxylipins we measured even after heat shock treatment (Fig. 1), which induces oxylipin accumulation in the charophyte alga *K. nitens* (Monte *et al*.,2020). Conversely, we detected OPDA and dn-OPDA in all streptophyte species (Fig. 1, S1, S2). It is worth mentioning that the isomers dn-*cis*-OPDA and dn-*trans*-OPDA appear as one single peak, that we collectively termed dn-OPDA, in contrast to the dn-*iso*-OPDA isomer, which could be clearly differentiated (Monte et al., 2018). Our results suggest that OPDA and dn-OPDA biosynthesis emerged in charophytes and is conserved in land plants. These results are consistent with previous detection of these two molecules in *K. nitens, M. polymorpha, P. patens, H. selago*, and *A. thaliana* (Koeduka et al., 2015; Chini et al., 2018; Monte et al., 2018, 2020, 2022; Mukhtarova et al., 2020). Consistently, we found putative orthologous AOS sequences in all analysed streptophytes lineages but not in chlorophytes, whereas AOC orthologs were exclusively present in land plants (Fig. 2; Supplementary Table S1). Sequence homology search in additional sequenced charophytes revealed that only *K. nitens* and *Spirogloea muscicola* possess AOS orthologs, while several charophyte algae contain OPR1/OPR2 orthologs and members of the GH3 family distinct from JAR1 (Supplementary Table S1). This indicates that OPDA/dn-OPDA biosynthesis occurs in streptophytes (Fig. 1, 2) and supports that AOC is not strictly necessary for OPDA synthesis, as previously proposed (Hamberg & Fahlstadius, 1990; Grechkin & Hamberg, 2000; Medvedeva et al., 2007). Moreover, the lack of AOS orthologs in some charophytes suggests that OPDA/dn-OPDA biosynthesis may have been lost in some extant charophyte clades.

**Figure 1:**
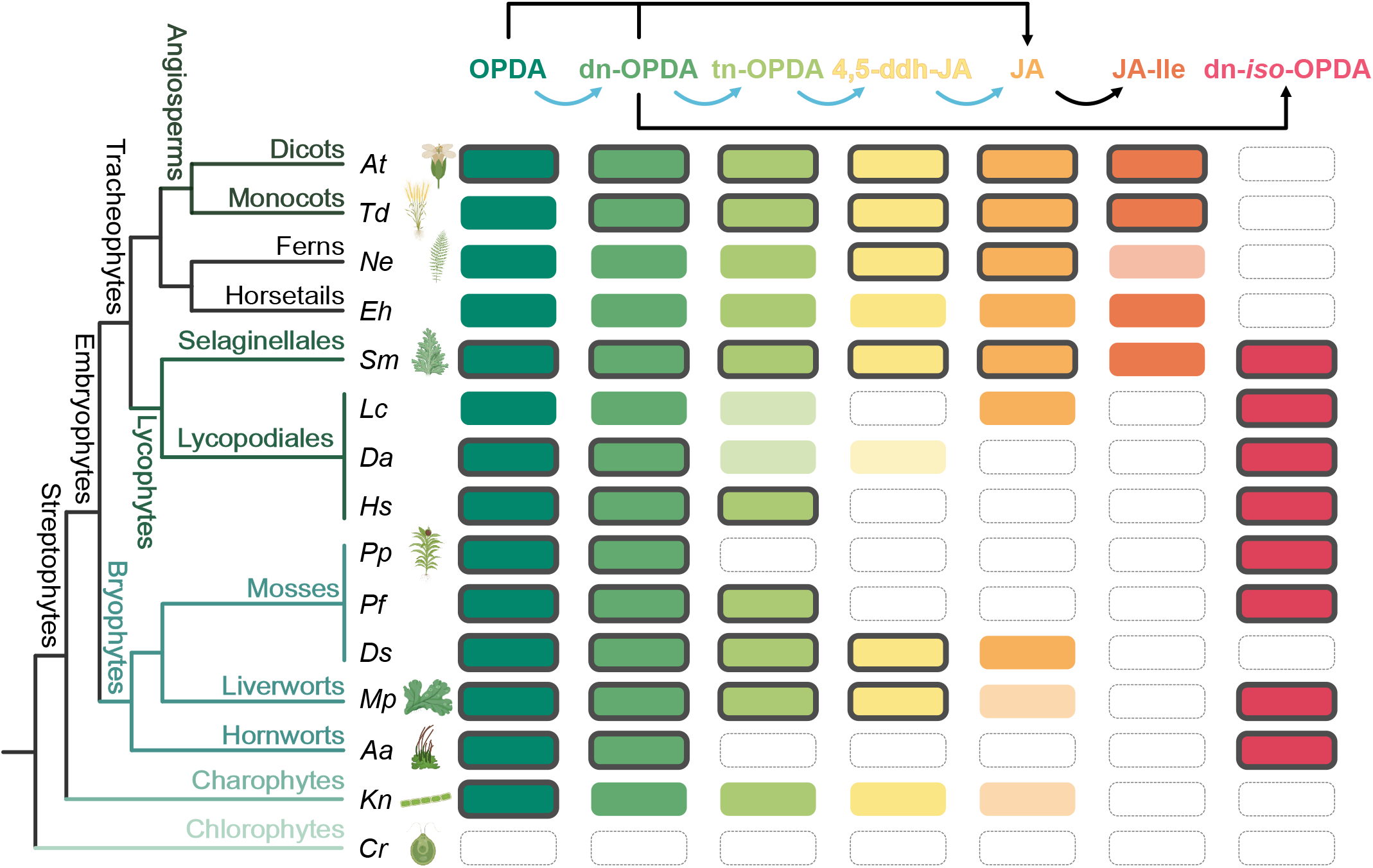
Jasmonates profile in the green lineage. Summary of jasmonates accumulation in representative species from the green lineage. Top, schematic representation of jasmonates biosynthesis where blue arrow indicates OPR3-independent synthesis of JA; left, phylogeny of the analyzed species. Solid colored rectangle, presence of the corresponding jasmonate; light colored rectangle, concentration close to the detection limit; grey border, jasmonates accumulation rapidly induced by wounding in land plants or heat shock in algae. At, *Arabidopsis thaliana;* Td, *Triticum durum* (wheat); Ne, *Nephrolepis exaltata*; Eh, *Equisetum hyemale*; Sm, *Selaginella moellendorffii*; Lc, *Lycopodium clavatum*; Da, *Diphasiastrum alpinum;*Hs, *Huperzia selago;* Pp, *Physcomitrium patens*; Pf, *Polytrichastrum formosum;* Ds, *Dicranum scoparium*; Mp, *Marchantia polymorpha*; Aa, *Anthoceros agrestis*; Kn, *Klebsormidium nitens*; Cr, *Chlamydomonas reinhardtii*.

**Figure 2:**
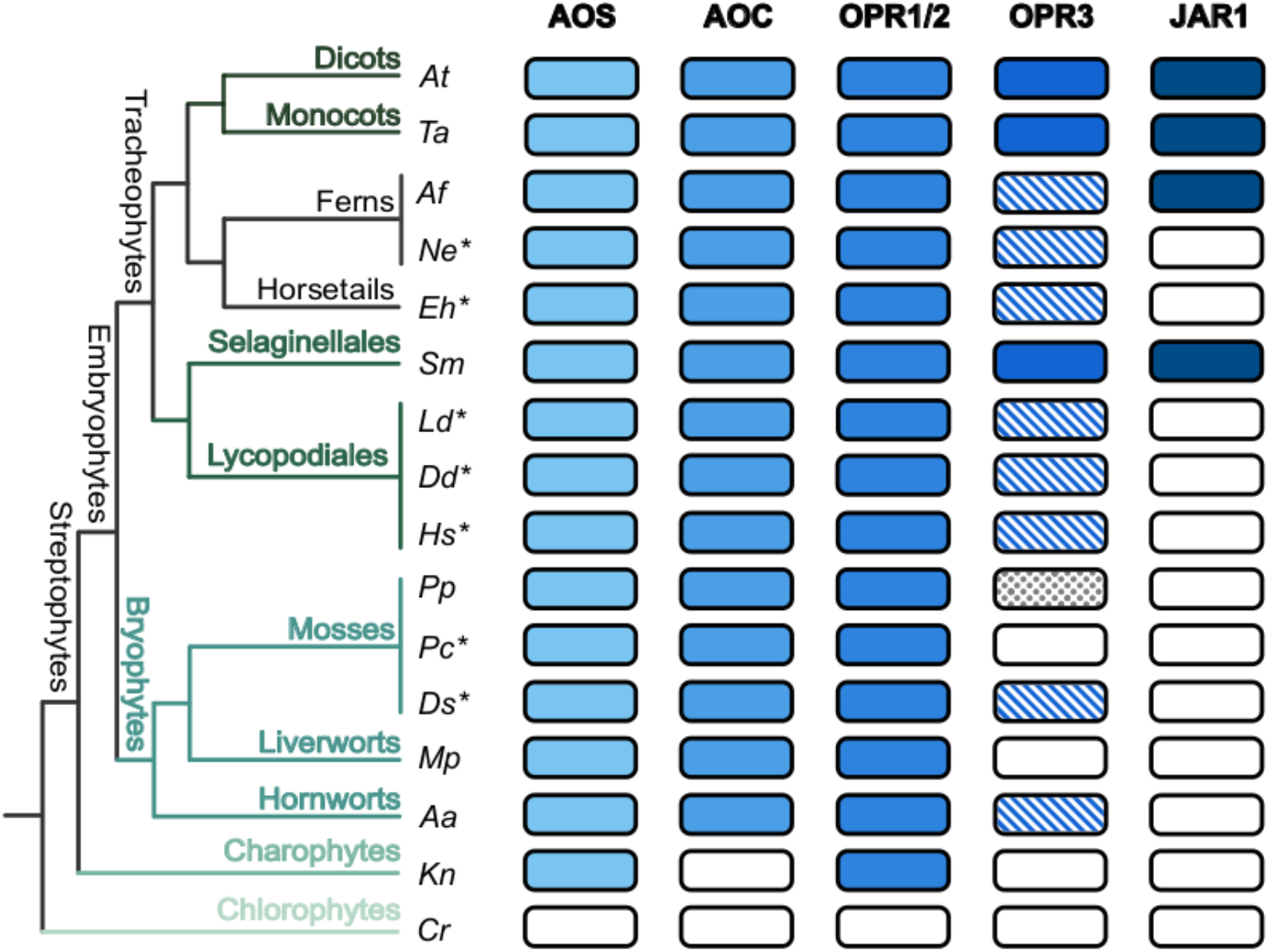
Conservation of jasmonates biosynthetic enzymes in the green lineage. Left, phylogeny of the analyzed species; asterisks indicate transcriptomes. Colored rectangle, presence of the corresponding enzyme; diagonal stripes in OPR3, sequences whose OPR3 function has not been tested; grey dots in PpOPR3, enzyme without OPR3 activity. At, *Arabidopsis thaliana*; Ta, *Triticum aestivum* (wheat); Af, *Azolla filiculoides*; Ne, *Nephrolepis exaltata*; Eh, *Equisetum hyemale*; Sm, *S. moellendorffii*; Ld, *Lycopodium deuterodensum*; Dd, *Diphasiastrum digitatum*; Hs, *Huperzia selago*; Pp, *Physcomitrium patens*; Pc, *Polytrichum commune*; Ds, *Dicranum scoparium*; Mp, *Marchantia polymorpha*; Aa, *Anthoceros agrestis*; Kn, *Klebsormidium nitens*; Cr, *Chlamydomonas reinhardtii*.

We observed an increase in OPDA accumulation after heat shock in *K. nitens* and after wounding in most land plants except one lycophyte, the horsetail *E. hyemale*, the fern *N. exaltata* and wheat (Fig. 1, S1). The stress-induced accumulation of dn-OPDA was not observed in the charophyte *K. nitens* and the abovementioned lycophyte, fern, and horsetail (Fig. 1, S2). Nevertheless, the horsetail *E. hyemale* exhibited elevated levels of OPDA and dn-OPDA already in control conditions, probably indicative of already stressed plants. Thus, we confirmed that OPDA/dn-OPDA biosynthesis emerged in streptophyte algae and is conserved across land plants.

### The ancestral OPR1/OPR2-dependent pathway produces JA and predates OPR3 evolution

OPDA and dn-*cis*-OPDA are precursors of JA through the canonical OPR3-dependent biosynthetic pathway starting with the reduction by OPR3 and subsequent beta-oxidation cycles (Wasternack & Hause, 2013). The discovery of an alternative OPR3-independent pathway in *A. thaliana* showed that OPDA can undergo successive cycles of beta-oxidation giving rise to dn-*cis*-OPDA, tn-OPDA and 4,5-ddh-JA, which is then reduced to JA by the cytosolic OPR1 and OPR2 (Chini *et al*., 2018). To identify whether the OPR3-independent pathway operates in plant lineages other than dicots, we first monitored the accumulation of the specific derivatives of the OPR1/OPR2 pathway tn-OPDA and 4,5-ddh-JA in the different species. These two molecules were detected in representatives of all streptophyte lineages except in hornworts and some mosses and lycophytes (Fig. 1, S3, S4). Furthermore, we detected the concurrent presence of 4,5-ddh-JA and JA in almost all species synthesizing 4,5-ddh-JA, suggesting that 4,5-ddh-JA might be generally reduced to JA by OPR1/OPR2 orthologs in streptophytes (Fig. 1, S4, S5). Consistently, our phylogenetic analyses revealed that class I OPRs (OPR1/OPR2) are present in all streptophytes lineages (Fig. 1, S6, Table S1), indicating that class I OPRs might have evolved in the common ancestor of all streptophytes. In the case of class II OPRs (OPR3-like) our phylogenetic analysis identified homologous sequences in land plants but not in charophyte algae, indicating that OPR3 function evolved later than OPR1/OPR2. Thus, the OPR1/OPR2-dependent JA biosynthetic pathway evolutionary precedes the OPR3-dependent pathway.

### JA-Ile biosynthesis emerged within lycophytes

JA accumulation and emergence of JAR1, the enzyme conjugating JA to Ile, were required for the development of JA-Ile biosynthesis (Staswick & Tiryaki, 2004). We observed that *K. nitens*, bryophytes, and Lycopodiales (a group of lycophytes) showed reduced levels of JA, sometimes close to the detection limit or even undetectable, whereas vascular plants such as the lycophyte *S. moellendorffii*, ferns, horsetails, and angiosperms exhibited elevated JA levels (Fig. 1, S5).

Phylogenetic analysis of GH3 proteins defined a clade that only contains proteins with proven JAR1 activity, spermatophyte and monilophyte sequences and one protein from the lycophyte *S. moellendorffii* (SmJAR1), which has been experimentally shown to have JAR1 activity (Pratiwi et al., 2017) (Fig. 2, S7, Table S1). Consistently, JA-Ile measurements only detected JA-Ile in angiosperms and monilophytes (ferns and horsetails), but we did not observe JA-Ile accumulation upon wounding after 5 or 30 min in any lycophyte (Fig. 1, 3). This result challenges the previous assumption that JA-Ile is present in all vascular plants based on the detection of JA-Ile in *S. moellendorffii* (Pratiwi et al., 2017). Further analyses showed that synthesis of JA-Ile in wounded *S. moellendorffii* only occurred at very late time points (3 and 6 hours) compared to the quick accumulation of dn-*iso*-OPDA within 5 minutes after wounding (Fig. 3, 4, S8). This result suggests that JA-Ile might not act as a stress response hormone in this plant. Thus, even though increased JA levels and accumulation of JA-Ile in *S. moellendorffii* point towards the emergence of JA-Ile biosynthesis in lycophytes, the role of JA-Ile as the main hormone mediating the response to wounding might have been acquired later on in evolution.

**Figure 3:**
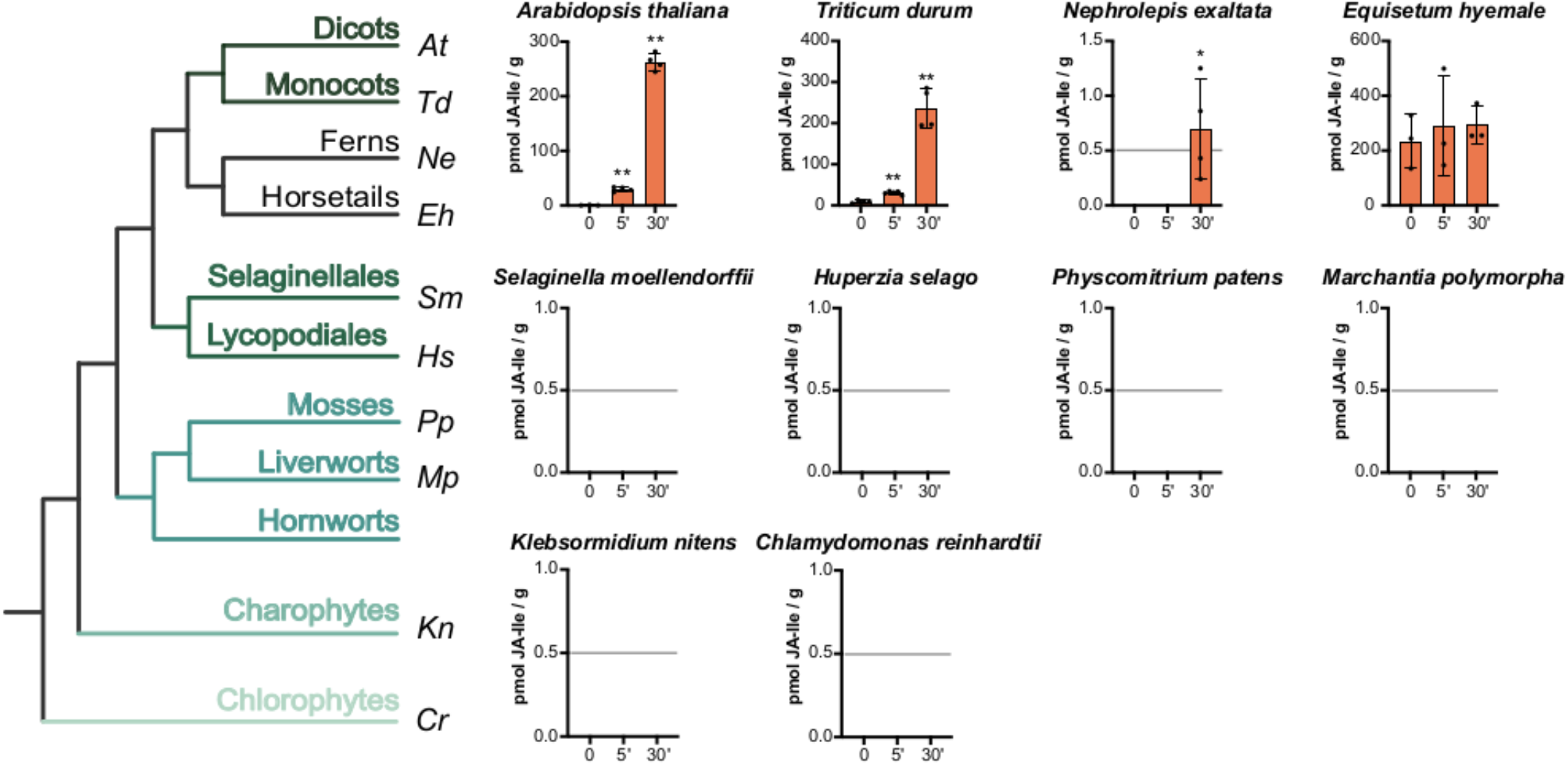
JA-Ile accumulates in angiosperms and monilophytes. Time-course accumulation of JA-Ile [pmoles/fresh weight (g)] in response to wounding in land plants or heat shock in algae. Data shown as mean ± s.d. of three or four biological replicates. Experiments were repeated 2 to 4 times with similar results. Asterisks indicate significant differences between wounded and unwounded samples according to a Student’s t-test analysis (* p-val < 0.05; ** p-val < 0.01). Horizontal grey line shows the detection limit (0.5 pmol). Left, phylogeny of the analyzed species.

**Figure 4:**
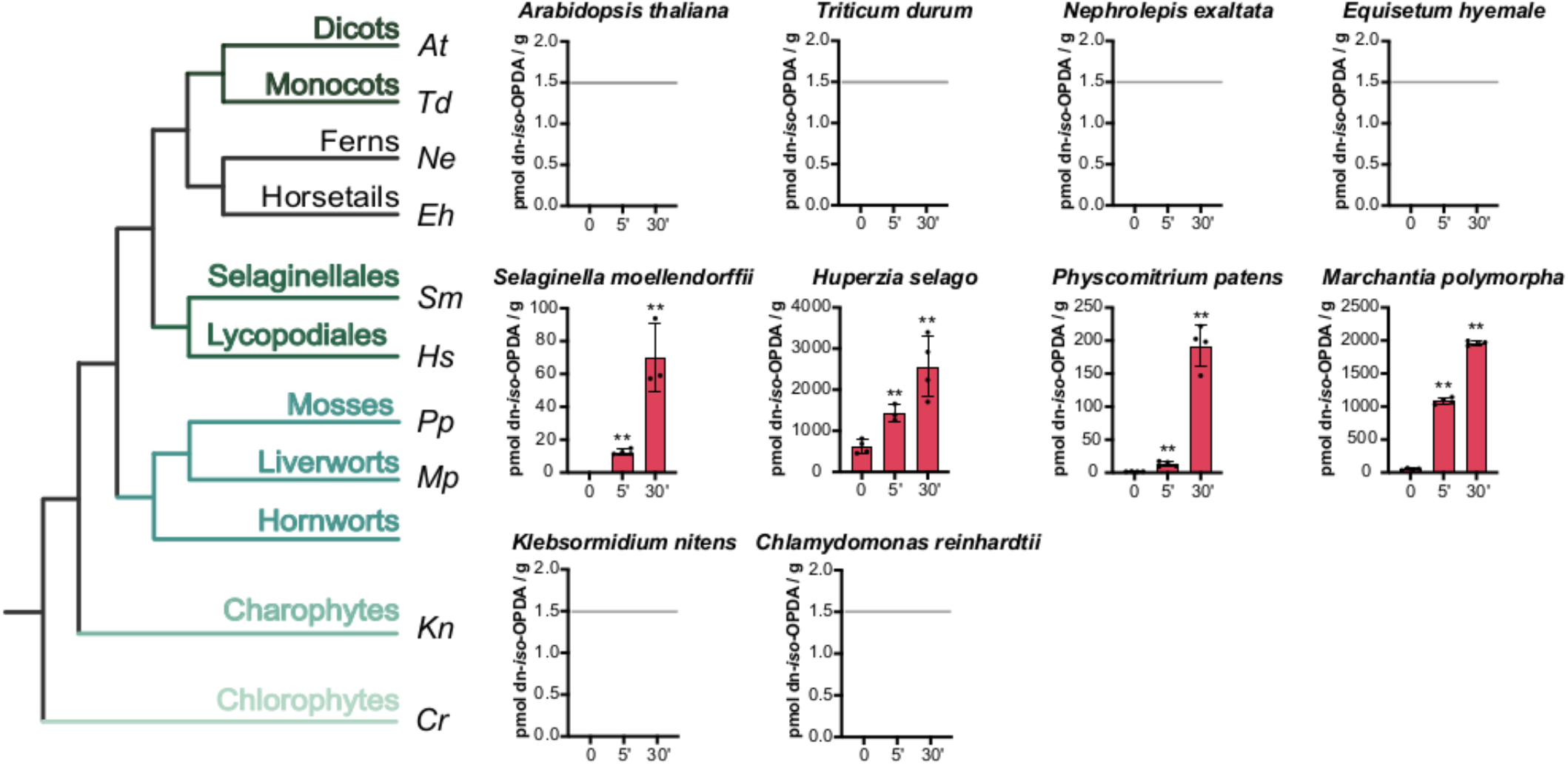
Lycophytes and bryophytes accumulate dn-*iso*-OPDA upon wounding. Time-course accumulation of dn-*iso*-OPDA [pmoles/fresh weight (g)] in response to wounding in land plants or heat shock in algae. Data shown as mean ± s.d. of three or four biological replicates. Experiments were repeated 2 to 4 times with similar results. Asterisks indicate significant differences between wounded and unwounded samples according to a Student’s t-test analysis (** p-val < 0.01). Horizontal grey line shows the detection limit (1.5 pmol). Left, phylogeny of the analyzed species.

### Dn-*iso*-OPDA might act as a hormone beyond bryophytes

Dn-*iso*-OPDA and its isomer dn-*cis*-OPDA act as COI1/JAZ ligands in *M. polymorpha* (Monte et al., 2018). To explore whether these two hormones were conserved across or beyond bryophytes, we measured dn-*iso*-OPDA in different species. Consistent with our previous results in *K. nitens* (Monte *et al*., 2020), we did not detect dn-*iso*-OPDA in this charophyte, even after heat shock treatment (Fig. 1, 4). Notably, all bryophytes and lycophytes except the moss *D. scoparium* synthesized both dn-*cis*-OPDA and dn-*iso*-OPDA (Fig. 1, 4, S2), indicating that these two molecules could exert the same function in bryophytes and lycophytes beyond *M. polymorpha*. Recently, we also defined Δ^4^-dn-OPDA, a dn-OPDA-like molecule, as an additional ligand of the jasmonate COI1/JAZ receptor in *M. polymorpha* (Kneeshaw *et al*., 2022). Our oxylipin analyses showed that only two bryophytes, *M. polymorpha* and *P. formosum*, accumulated Δ^4^-dn-OPDA (Fig. S9). This indicates that Δ^4^-dn-OPDA might be relevant in a limited number of plant species, in contrast to the broader presence of dn-OPDA isomers and JA-Ile across different lineages. Unlike the other bryophytes, *D. scoparium* accumulated 4,5-ddh-JA and JA (Fig. 1), suggesting a shift in the dn-*cis*-OPDA fate towards beta-oxidation rather than isomerization to dn-*iso*-OPDA. Nevertheless, we detected 3,7-ddh-JA in *D. scoparium* (Fig. S10), which is the putative beta-oxidation derivative of dn-*iso*-OPDA, indicating that *D. scoparium* might as well produce dn-*iso*-OPDA that is quickly processed. Alternatively, 3,7-ddh-JA (*iso* form) could be an isomerization product of 4,5-ddh-JA (*cis* form). Besides being a dn-*cis*-OPDA derivative, dn-*iso*-OPDA can be also a product of *iso*-OPDA through one cycle of beta-oxidation (Mukhtarova *et al*., 2020). Thus, we measured *iso*-OPDA and found an elevated concentration of this oxylipin in all mosses and lycophytes, and minute amounts after wounding in the hornwort *A. agrestis* (Fig. 1, S11). Wounding increased the concentration of *iso*-OPDA in these plants but decreased the concentration of *iso*-OPDA in *A. thaliana* (Fig. 1, S10). These results indicate that the source of dn-*iso*-OPDA is different between liverworts, and mosses, lycophytes and maybe hornworts.

One of the most striking results of our oxylipin profile analysis was the negative correlation between JA-Ile and dn-*iso*-OPDA in all land plants species except *S. moellendorffii*, which indicates a dichotomy in OPDA/dn-*cis*-OPDA fate towards either dn-*iso*-OPDA or JA-Ile (Fig. 1, 3, 4). The mutually exclusive presence of dn-*iso*-OPDA and JA-Ile and their specific role as COI1/JAZ ligands suggest that dn-*iso*-OPDA and JA-Ile might be equivalent hormones perceived by COI1/JAZ in different plant lineages. The identification of dn-*iso*-OPDA in lycophytes, where only *S. moellendorffii* accumulates JA-Ile at very late time points after wounding (Fig. 1, 3, 4, S8), suggests that instead of JA-Ile, lycophytes and bryophytes employ dn-*iso*-OPDA as the main COI1/JAZ ligand.

## Discussion

Distinct jasmonates activate independent but conserved signaling pathways in plants. OPDA and dn-*cis*-OPDA play a COI1-independent role in thermotolerance in streptophytes (Monte *et al*., 2020), whereas dn-*cis*-OPDA, dn-*iso*-OPDA and Δ^4^-dn-*iso*-OPDA in *M. polymorpha*, and JA-Ile in angiosperms activate the COI1-dependent pathway (Fonseca et al., 2009; Sheard et al., 2010; Monte et al., 2018). In this work, we addressed how widespread these bioactive jasmonates are in representative plant lineages and how their biosynthetic pathways evolved. In addition to several species (*K. nitens*, bryophytes, some lycophytes, and angiosperms) where jasmonate profiles had already been studied, here we analyzed chlorophytes, hornworts, additional bryophytes, lycophytes, ferns, and horsetails. Our phylogenetic analyses of the key enzymes for jasmonate biosynthesis showed the presence of putative orthologs of AOS and OPR1/OPR2 in charophytes and their conservation in all land plants. Consistently, we detected OPDA and dn-*cis*-OPDA in *K. nitens* and land plants, but not in the chlorophyte *C. reinhardtii*, which support the idea that these molecules play conserved COI1-independent role in thermotolerance in streptophytes (Monte *et al*., 2020). The absence of orthologs for the biosynthetic enzymes AOS and AOC in the charophyte *Chara braunii* suggests that OPDA/dn-*cis*-OPDA synthesis was lost in some charophyte lineages (Nishiyama *et al*., 2018), yet the related species *C. australis* has been reported to synthesize JA (Beilby *et al*., 2015). AOC is not strictly required for OPDA/dn-OPDA biosynthesis (Hamberg & Fahlstadius, 1990; Grechkin & Hamberg, 2000; Medvedeva *et al*., 2007), as shown in *K. nitens*,which lacks AOC orthologs but still synthesizes OPDA and dn-*cis*-OPDA. Thus, the presence of AOS in *S. muscicola* suggests that this Zygnematophyceae alga, considered to belong to the sister lineage of land plants, is also capable of synthesizing OPDA/dn-*cis*-OPDA.

Consistent with the well-established role of mechanical wounding in jasmonates accumulation, we observed an increase in jasmonates accumulation upon wounding in all land plant species except for the horsetail *E. hyemale* (Fig. 1). Nevertheless, we detected different wound-induced accumulation patterns depending on the species and individual jasmonates. Differences in plant morphology including coriaceous leaves or fibrous stems co-related with attenuated response to wounding observed in the reduced JA-Ile accumulation in wounded *N. exaltata*. In contrast, the horsetail *E. hyemale* contained elevated levels of all jasmonates prior to wounding, which might indicate that the plants were already stressed in basal conditions. We observed a remarkable accumulation of 4,5-ddh-JA at every time point in *E. hyemale*, confirming the previously reported detection of this metabolite in other *Equisetum* species (Dathe *et al*., 1989).

OPDA has been long known as a precursor of JA-Ile in angiosperms. Indeed, OPDA can be converted to JA through two different pathways: the canonical pathway involving the peroxisomal enzyme OPR3 and several cycles of beta-oxidation, or the alternative pathway starting with the beta-oxidation (and production of dn-*cis*-OPDA) leading to the transformation of 4,5-ddh-JA into JA through cytosolic OPR1/OPR2 enzymes (Chini *et al*., 2018). Concurrent presence of 4,5-ddh-JA and JA in all streptophyte lineages together with the identification of OPR1/OPR2 orthologs in all streptophytes and the lack of OPR3 orthologs in charophytes, indicate that the alternative biosynthetic pathway involving OPR1/OPR2 is ancestral and evolutionarily precedes the OPR3-dependent pathway. Regarding the evolution of the OPR3 pathway, our phylogenetic analysis showed putative orthologs in several land plants. For instance, the *P. patens* OPR3a exhibits the canonical FH residues of AtOPR3 yet lacks OPR3 activity (Bandara et al., 2012). Conversely, SmOPR5 shows divergent residues (WH) but exhibits OPR3 function (Pratiwi et al., 2017). Thus, phylogenetic analyses should be complemented with functional characterization of OPR enzymes to define when OPR3 acquired its function on OPDA reduction during evolution. Nevertheless, considering the lower JA levels detected in charophytes, bryophytes and Lycopodiales, compared to monilophytes and angiosperms, and the demonstrated OPR3 activity of SmOPR5, JA biosynthesis by OPR3 enzymes may have appeared in vascular plants as a result of neo-functionalization of class II OPR enzymes. The emergence of OPR3 function, showing stereoselectivity for OPDA and localization in the peroxisome, likely increased the JA conversion rate by maintaining the enzyme OPR3 and its substrate OPDA in the same subcellular compartment, the peroxisome, and therefore was crucial for the evolution of JA-Ile biosynthesis. Prevention of OPDA release from the peroxisome might have limited the activation of COI1-independent effects due to OPDA properties as reactive electrophilic species (Monte *et al*., 2020). Complementary metabolomic analysis of the OPR3 derivatives OPC-8, OPC-6 and OPC-4 could not be conducted due to the fast turn-over of these molecules (Schulze *et al*., 2019).

Evolution of JAR1 activity within the GH3 family was sufficient to give rise to JA-Ile in lycophytes. Notably, we only detected JA-Ile in *S. moellendorffii* but not in lycophytes belonging to Lycopodiales, consistent with the absence of JAR1 orthologs in their transcriptomes and previous oxylipin measurements in *H. selago* (Monte *et al*., 2022). Nevertheless, OPR3 and JAR1-like sequences were identified in the genome of *Isoëtes taiwanensis*, belonging to the third group of lycophytes, the Isoetales, and therefore similar to Selaginellales. Based on these analyses *I. taiwanensis* might be able to synthesize JA-Ile. Genome sequencing and jasmonate profiling of additional lycophytes will be instrumental to clarify the origin and conservation of JA-Ile biosynthetic enzymes in this lineage. Functional analysis of SmOPR5 and SmJAR1 confirmed their OPR3 and JAR1 activity, despite showing divergence in key residues present in their *A. thaliana* counterparts (Pratiwi et al., 2017, Fig S6, Table S1). We showed that wounding of *S. moellendorffii* induced JA-Ile accumulation much later than the quick accumulation of dn-*iso*-OPDA (5 minutes vs 3 hours), which questions the JA-Ile role as stress hormone in *Selaginella*. Nonetheless, JA-Ile might still regulate different processes in *S. moellendorffii*.

Instead of JA-Ile, the COI1/JAZ ligands in the liverwort *M. polymorpha* are two isomers of dn-OPDA (dn-*cis*-OPDA and dn-*iso*-OPDA) and Δ^4^-dn-*iso*-OPDA (Monte et al., 2018; Kneeshaw et al., 2022). Dn-*cis*-OPDA can be synthesized from OPDA through one cycle of beta-oxidation, but the main source of dn-*cis*-OPDA in *M. polymorpha* is the fatty acid 16:3 (Monte et al., 2018; Soriano et al., 2022). Dn-*cis*-OPDA can isomerize into dn-*iso*-OPDA by a yet unknown mechanism that might involve an enzymatic reaction (Monte et al., 2018). In the moss *P. patens*, OPDA can undergo a similar isomerization to form *iso*-OPDA, that in turn can give rise to dn-*iso*-OPDA through beta-oxidation (Mukhtarova *et al*., 2020). We detected *iso*-OPDA in all mosses, the hornwort, and the lycophytes, but not in the charophyte *K. nitens* or in *M. polymorpha*, indicating that the synthesis of the *iso* forms is specific to land plants and that *iso*-OPDA is potentially a precursor of dn-*iso*-OPDA in mosses, hornworts, and lycophytes but not in liverworts. Conversion of OPDA to *iso*-OPDA was identified in the gut of herbivore insects as a detoxification mechanism to inactivate OPDA (Dabrowska *et al*., 2009), but the responsible enzyme has no orthologs in plants (Dabrowska & Boland, 2007). Our identification of plant lineages synthesizing *iso*-OPDA and dn-*iso*-OPDA could represent a first step to identify this putative plant isomerase.

The identification of dn-OPDA-related molecules as COI1/JAZ ligands in *M. polymorpha* (Monte et al., 2018) and the proposed emergence of JA-Ile biosynthesis in vascular plants (Pratiwi et al., 2017) anticipated that additional plant lineages could use dn-OPDAs as hormones. Our oxylipin profiling revealed that dn-*cis*-OPDA is produced in the charophyte *K. nitens* and all land plants, whereas dn-*iso*-OPDA accumulated in bryophytes and lycophytes. Notably, Δ^4^-dn-*iso*-OPDA seems to be exclusively in setaphyta (mosses + liverworts). The hypothesis that other bryophytes in addition to *M. polymorpha* use dn-OPDA relatives as the hormones perceived by COI1/JAZ is consistent with the growth inhibition by OPDA in the moss *P. patens* and the hornwort *A. agrestis* (Monte et al., 2018), which is likely due to OPDA conversion to dn-*cis*-OPDA and dn-*iso*-OPDA. Moreover, these bryophytes were insensitive to JA or JA-Ile, in keeping with conserved COI1 sequences lacking a key alanine residue in their binding pocket that cannot accommodate a larger ligand like JA-Ile, similar to *M. polymorpha* (Monte et al., 2018). Our results support that JA-Ile is not a hormone in bryophytes as they do not synthesize nor perceive this molecule. Notably, we recently discovered that JAZs also contribute to ligand specificity, together with COI1 (Monte *et al*., 2022). Thus, the COI1/JAZ co-receptor of the lycophyte *H. selago*, which accumulates both dn-*cis*-OPDA and dn-*iso*-OPDA but not JA-Ile, is able to bind all three molecules due to specific amino acid combinations where the HsCOI1 protein is similar to angiosperms COI1 and the HsJAZ is similar to bryophyte JAZs (Monte *et al*., 2022). Biosynthesis of dn-*iso*-OPDA in bryophytes and lycophytes indicate that these two plant lineages differ from all other vascular plants.

Notably, we identified a negative co-relation between dn-*iso*-OPDA and JA-Ile in all plants except in *S. moellendorffii*, which is capable of synthesizing both molecules. Thus, we hypothesized that dn-*iso*-OPDA acts as a stress hormone perceived by COI1/JAZ in lycophytes as well as in bryophytes, as we experimentally demonstrated in *M. polymorpha* and *H. selago* (Monte et al., 2018; Monte et al., 2022)Monte et al., 2022. The very late accumulation of JA-Ile in wounded *S. moellendorffii* likely indicates that JA-Ile does not act as a rapidly induced stress hormone. Genetic dissection of JA-Ile biosynthesis and signaling in *S. moellendorffii* and other lycophytes will reveal whether JA-Ile plays a similar role to other vascular plants beyond wounding. Functional analyses in lycophytes belonging to Isoetales will be essential to decipher whether they synthesize and/or use JA-Ile as a signaling molecule. The absence of JA-Ile in several Lycopodiales and the accumulation of dn-*iso*-OPDA indicate that this group of lycophytes likely use dn-*iso*-OPDA as a stress hormone, similar to *H. selago*, thus sharing some features with bryophytes. The COI1/JAZ co-receptor complex from *H. selago* and other lycophytes is able to perceive JA-Ile in addition to dn-OPDA isomers (Monte *et al*., 2022), which is particularly relevant in the case of lycophytes that synthesize both types of molecules, such as *S. moellendorffii* and possibly *Isoëtes sp*. How different ligands employ the same co-receptor complex in lycophytes to regulate stress responses or development represent exciting future questions. Our previous work showed dn-*cis*-OPDA and dn-*iso*-OPDA bind the COI1/JAZ receptor in *M. polymorpha*, but the reactive electrophilic nature of dn-*cis*-OPDA, similar to OPDA, provides additional COI1-independent effects protecting plants against heat shock (Monte *et al*., 2020). Uncontrolled release of reactive electrophilic species within the cell might be harmful and therefore must be tightly regulated. Based on our hormone measurements across the green lineage, we hypothesized that dn-*cis*-OPDA has two mutually exclusive fates converting to either dn-*iso*-OPDA in bryophytes and lycophytes or JA-Ile in the rest of vascular plants (and some lycophytes). Thus, dn-*iso*-OPDA and JA-Ile represent two analogous hormones devoid of reactive electrophilic properties that can specifically activate the jasmonate signaling pathway through COI1/JAZ binding in different plant lineages.

## Materials and Methods

### Plant material and growth conditions

Plants were grown in a growth chamber in short-day (10/14h photoperiod) at 21°C and 65% relative humidity as previously described (Chini et al., 2021). As spermatophytes representatives, *Arabidopsis thaliana* (Col-0 ecotype) and wheat *Triticum durum* (Ebel *et al*., 2018) plants were analysed in this study. 4-week-old soil-grown plants were wounded with forceps and leaves were collected 5 and 30 minutes after wounding. Leaves from 5/10 plants were collected per each sample and 4 samples were analysed per each time point. Unwounded plants (0) were included as control. Experiments were carried out at least 3 times with similar results. The monilophytes (*Nephrolepis exaltata* and *Equisetum hyemale*), lycophytes (*Selaginella moellendorffii*, *Huperzia selago*, *Diphasiastrum alpinum* and *Lycopodium clavatum*) and bryophytes (*Marchantia polymorpha*, *Physcomitrium patens*, *Dicranum scoparium*, *Polytrichastrum formosum* and *Anthoceros agrestis*) analysed in this study were also grown in soil in similar condition, at approximately 100% relative humidity. The bryophytes *Marchantia polymorpha*, *Physcomitrium patens* and *Anthoceros agrestis* were grown in GB5 plates. The wounding assays were carried out as described above. 3/4 samples were collected, and experiments were twice or three times with similar results. The algae *Klebsormidium nitens* (NIES-2285) and *Chlamydomonas reinhardtii* were grown at 21°C in a 16-h light/8-h dark cycle in GB5 and TAP medium respectively, as previously described (Monte *et al*., 2020). The algae were stressed at 37<u>°</u>C for one hour, and material was collected for jasmonate measurements after 1 hour. Unstressed algae (0) were included as control. 4 samples were collected, and experiments were repeated twice or three times with similar results.

### Gene identification

Sequences from representative plant lineages were retrieved using Arabidopsis AOS (AT5G42650), AOC1 (AT3G25760), OPR1 (AT1G76680), OPR2 (AT1G76690), OPR3 (AT2G06050) and GH3.11/JAR1 (AT2G46370) proteins as query in different databases: OneKP (https://db.cngb.org/onekp/), Phytozome (https://phytozome-next.jgi.doe.gov/), NCBI BLAST blastp suite (https://blast.ncbi.nlm.nih.gov/Blast.cgi), ORCAE (http://bioinformatics.psb.ugent.be/orcae/), MaizeGDB (https://maizegdb.org/), Wheat IWGSC (https://www.wheatgenome.org/), Spruce Genome Project (https://congenie.org/start), Fernbase (www.fernbase.org), MarpolBase (https://marchantia.info/) and Klebsormidium nitens genome project (http://www.plantmorphogenesis.bio.titech.ac.jp/~algae_genome_project/klebsormidium/index.html). In the absence of sequenced genomes/transcriptomes for some of the species used for jasmonates profiling, we employed related species as follows: *Polytrichum commune* instead of *P. formosum, L. deuterodensum* instead of *L. clavatum*,and *Diphasiastrum digitatum* instead of *D. alpinum*. Sequences from *T. aestivum* were included due to previous functional characterization of JAs biosynthetic azymes. *Azolla filiculoides* sequences were included to confirm the presence of JAR1 orthologs in ferns.

### Alignment and phylogenetic analyses

Protein alignments were obtained with MUSCLE (www.ebi.ac.uk/Tools/msa/muscle/), construction of phylogenetic tree was performed with PhyML (www.phylogeny.fr/index.cgi) and the Tree Viewer programme TreeDyn (www.phylogeny.fr/index.cgi) was used to generate the final tree as previously described (Chini *et al*., 2017).

### Chemicals

(-)-Jasmonic acid (JA), cis-12-oxo-phytodienoic acid (OPDA) and N-(-)-jasmonoyl isoleucine (JA-Ile) were purchased from OlChemim Ltd (Olomouc, Czech Republic), dinor-12-oxo-phytodienoic acid (dn-OPDA) from Cayman Chemical Company (Ann Arbor, MI, USA), whereas dn-*cis*-OPDA, dn-*iso*-OPDA, tn-OPDA, 4,5-ddh-JA and 3,7-ddh-JA were previously synthesized (Chini et al., 2018; Monte et al., 2018, 2020).

### Jasmonate measurements

Jasmonate measurements were performed as previously described (Chini *et al*., 2018; Monte *et al*., 2020) using biological samples of tissue pooled from 5-10 plants and at least three independent biological replicates were measured for each treatment. The experiment was repeated twice to four times with similar results. Data are shown as mean ± SD.

## Supporting information

Oxylipin evolution Suppl Figures bioRxiv

## Acknowledgements

This research was funded by the Spanish Ministry for Science and Innovation grant PID2019-107012RB-100 and “Severo Ochoa” Programme for Centres of Excellence SEV-2017-0712 funded by the MCIN/AEI/ 10.13039/501100011033. We thank Javier Martínez Abaigar and Encarnación Nuñez (University of La Rioja, Spain) for moss identification and providing *Huperzia selago* and *Lycopodium clavatum* plants, Begoña Benito (Polytechnic University of Madrid) for providing *Physcomitrium patens*, Vicente Mariscal (IBVF/CSIC, Seville) for providing *Chlamydomonas reinhardtii*, Jody Banks (Purdue University) for providing *Selaginella moellendorffii*, Marie-Genevieve Nicolas (Parc National des Ecrins, France), Paul Nicolas (University of Clermont-Ferrand, France), Anastasiia I Evkaikina and Olga Voitsekhovskaja (Komarov Botanical Institute, Russian Academy of Sciences, Russia) for kindly providing *Huperzia selago* plants.

## Author Contributions

R.S., A.C. and I.M. designed the research. A.C. and I.M. performed all experiments except the oxylipin measurements, performed by A.M.Z. and supervised by J.M.G-M. A.C., I.M. and R.S. analysed all data. A.C., I.M. and R.S wrote the manuscript. All authors revised and approved the final version of the manuscript.

