## Supplementary material for "Evolution of the jasmonate ligands and their biosynthetic pathways": Oxylipin evolution Suppl Figures bioRxiv

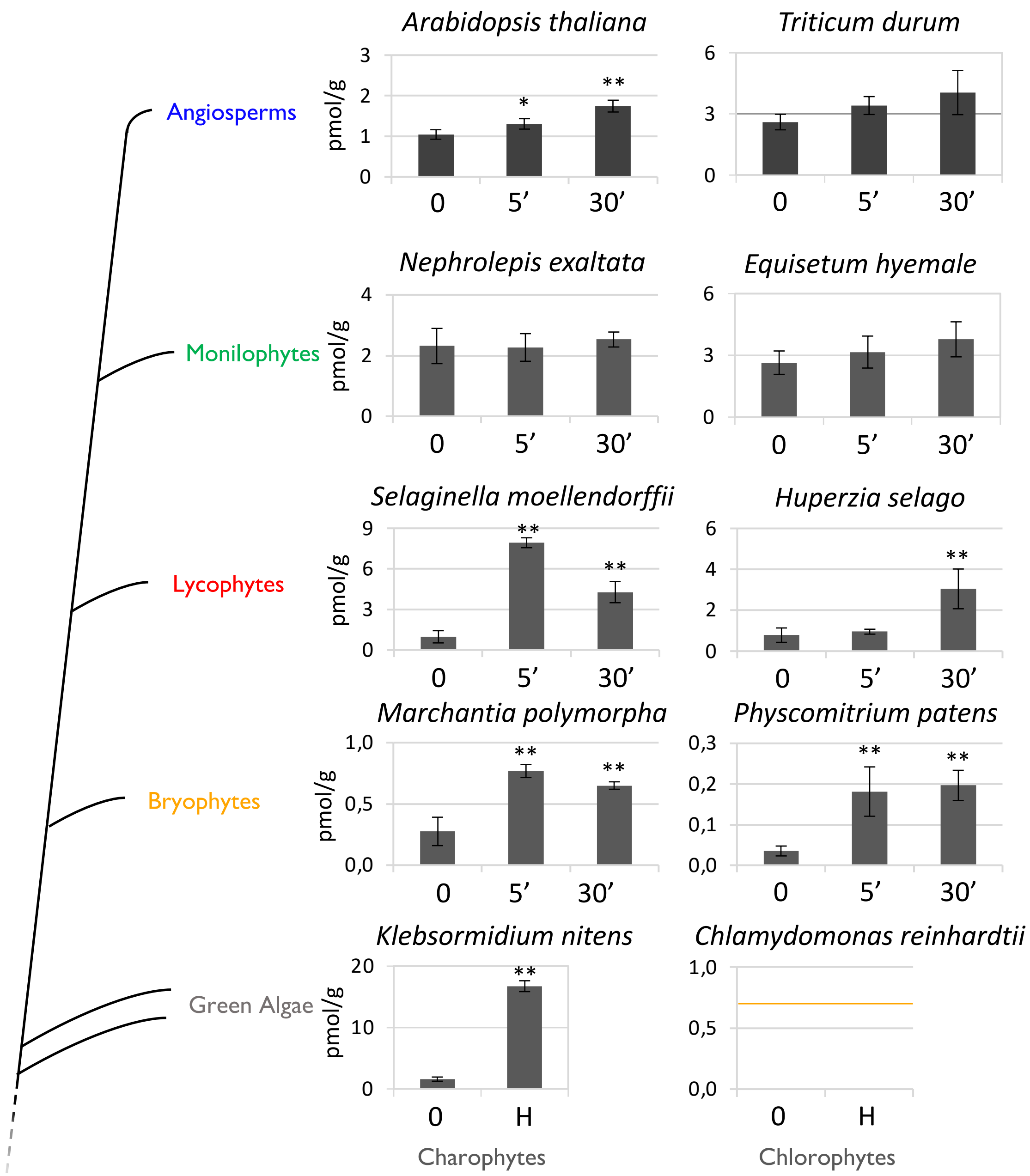


**Fig. S1. Accumulation of OPDA after wounding in representative plant species.**

Time-course accumulation of OPDA [nmoles/fresh weight (g)] in several plant species after wounding. Plants were wounded and damaged tissues were collected after the indicated times. Unwounded plants (0) were included as control. Algae were heat-stressed (H) for 1 hour at 37ºC. Data shown as mean ± s.d. of three or four biological replicates. Experiments were repeated 2 to 4 times with similar results. Asterisks indicate significant differences between wounded and unwounded samples according to a Student's t-test analysis (** p-val < 0.01). An orange line shows the detection limit (0.7 pmol) in the case of *Chlamydomonas reinhardtii*, in which OPDA was not detected. A schematic representation of a plant phylogenetic tree is also reported

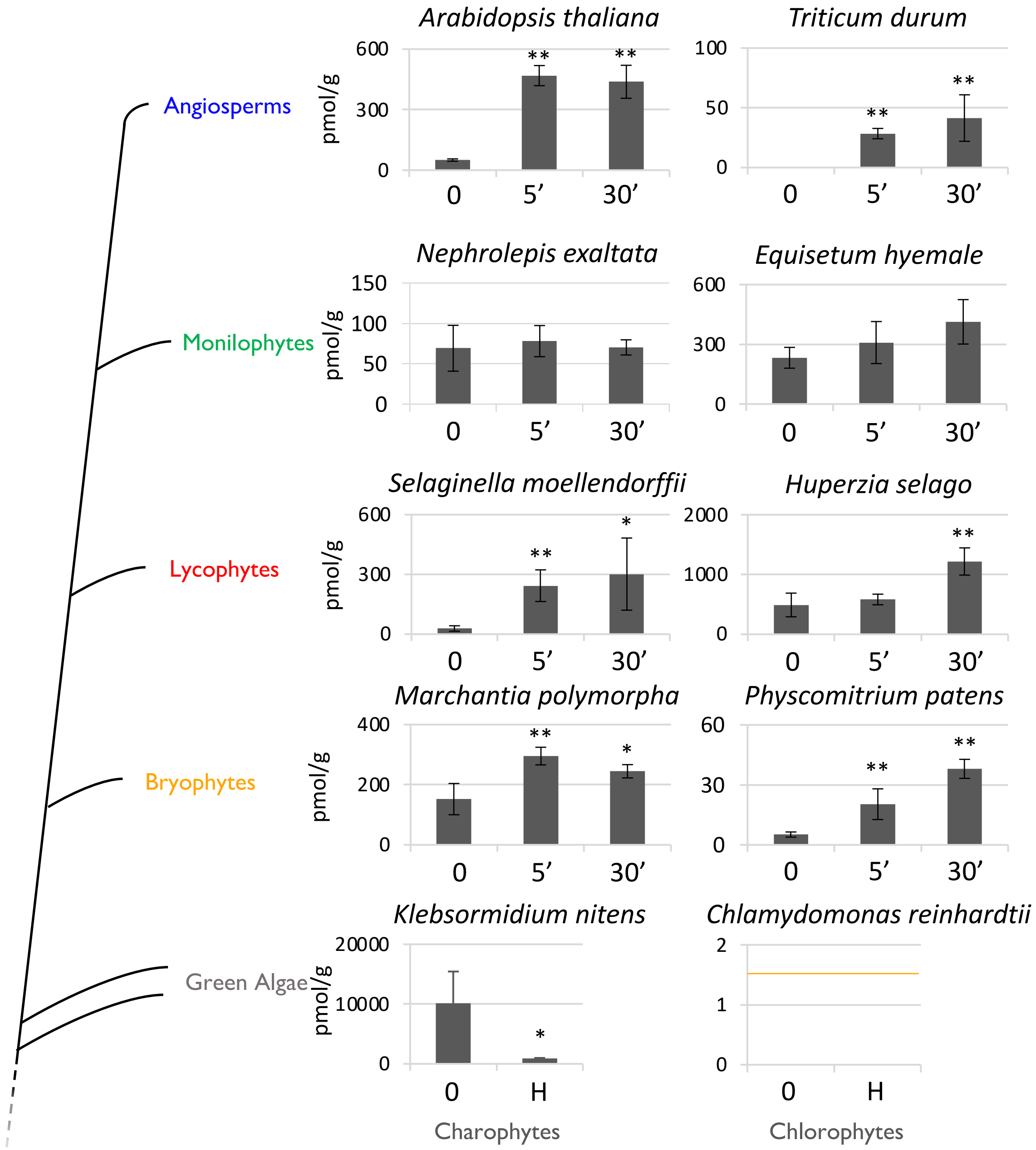


**Fig. S2. Accumulation of dn-OPDA after wounding in representative plant species**.

Time-course accumulation of dn-OPDA [pmoles/fresh weight (g)] after wounding in several plant species. Data shown as mean ± s.d. of three or four biological replicates. Experiments were repeated 2 to 4 times with similar results. Asterisks indicate significant differences between wounded and unwounded samples according to a Student's t-test analysis (* p-val < 0.5, ** p-val < 0.01). An orange line shows the detection limit (1.5 pmol) in the case of *Chlamydomonas reinhardtii*, in which dn-OPDA was not detected. On the left, a schematic representation of a plant phylogenetic tree is included.

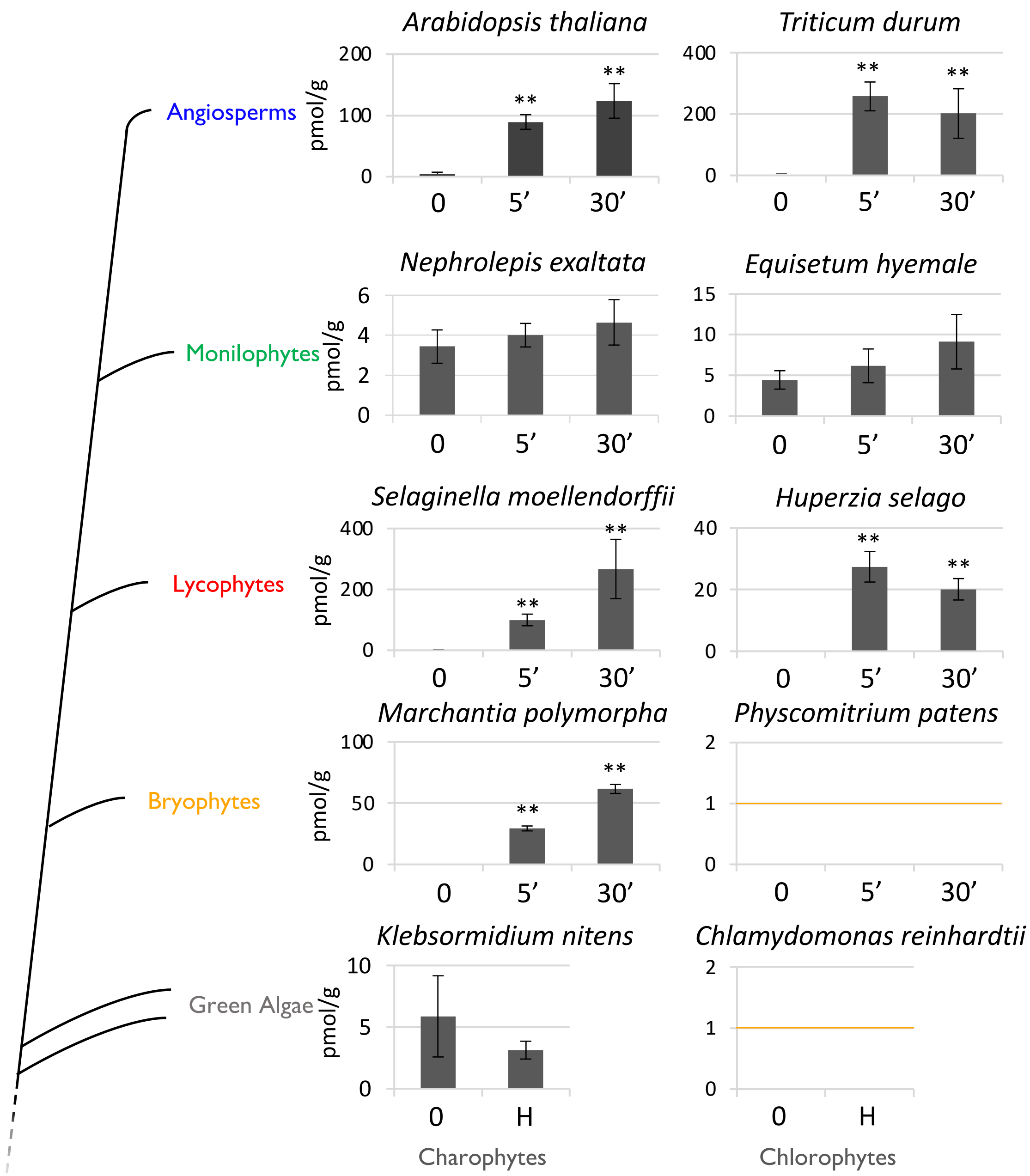


**Fig. S3. Accumulation of tn-OPDA after wounding in representative plant species**.

Time-course accumulation of tn-OPDA [pmoles/fresh weight (g)] after wounding in several plant species. Data shown as mean ± s.d. of three or four biological replicates. Experiments were repeated 2 to 4 times with similar results. Asterisks indicate significant differences between wounded and unwounded samples according to a Student's t-test analysis (** p-val < 0.01). An orange line shows the detection limit (1 pmol) in the case of plants in which OPDA was not detected. On the left, a schematic representation of a plant phylogenetic tree is included.

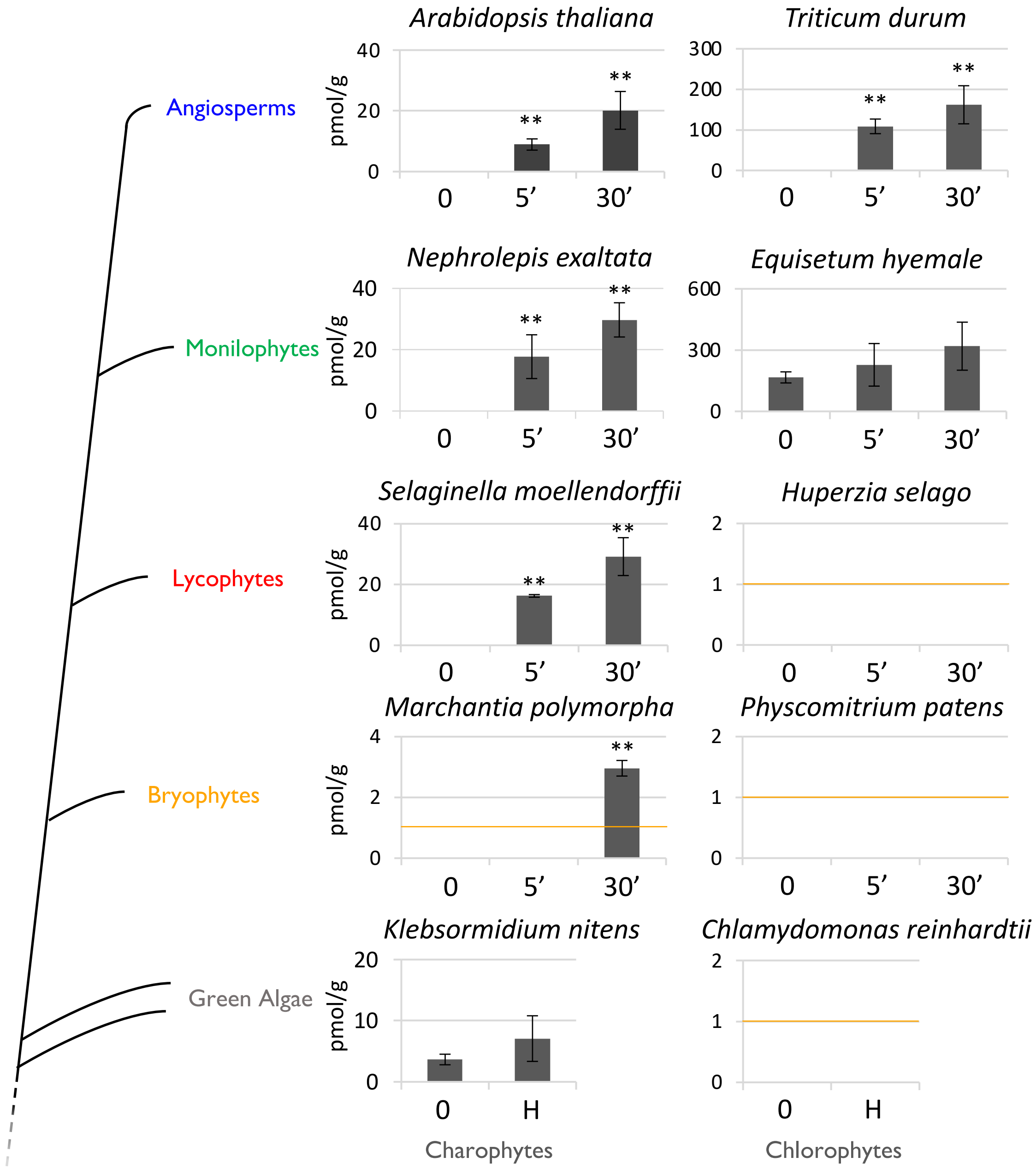


**Fig. S4. Accumulation of 4,5-ddh-JA after wounding in representative plant species**.

Time-course accumulation of 4,5-ddh-JA [pmoles/fresh weight (g)] in response to wounding in several plant species. Data shown as mean ± s.d. of three or four biological replicates. Experiments were repeated 2 to 4 times with similar results. Asterisks indicate significant differences between wounded and unwounded samples according to a Student's t-test analysis (** p-val < 0.01). An orange line shows the detection limit (1 pmol). A schematic representation of a plant phylogenetic tree is included on the left.

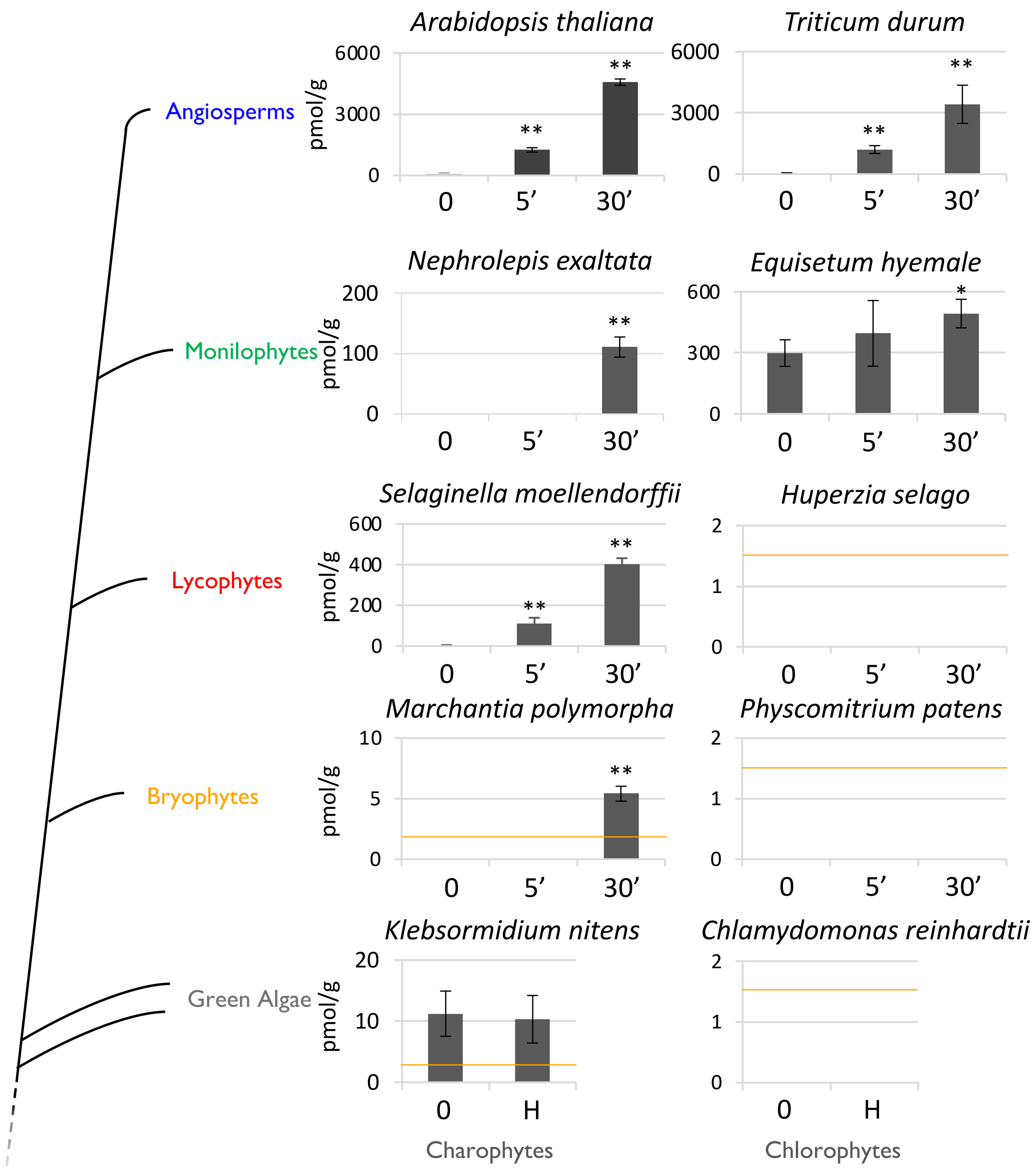


**Fig. S5. Accumulation of JA after wounding in representative plant species**.

Time-course accumulation of JA [pmoles/fresh weight (g)] after wounding in several plant species. Experiments were repeated 2 to 4 times with similar results. Data shown as mean ± s.d. of three or four biological replicates. Asterisks indicate significant differences between wounded and unwounded samples according to a Student's t-test analysis (** p-val < 0.01). An orange line shows the detection limit (1.5 pmol). On the left, a schematic representation of a plant phylogenetic tree is included.

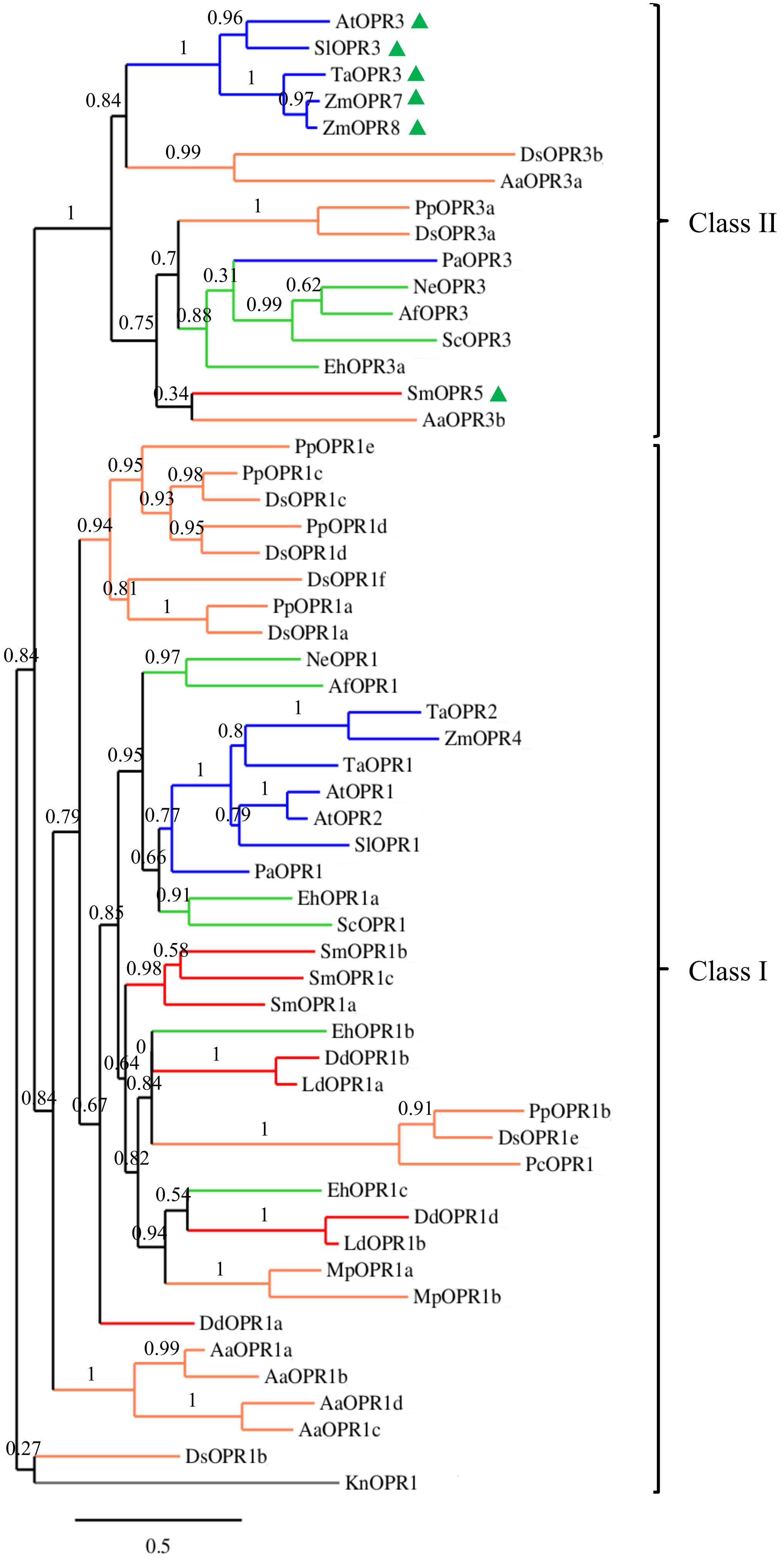
AtOPR3
SlOPR3
TaOPR3
ZmOPR7
ZmOPR8
DsOPR3b
AaOPR3a
PpOPR3a
DsOPR3a
PaOPR3
NeOPR3
AfOPR3
ScOPR3
EhOPR3a
SmOPR5
AaOPR3b
Class II
PpOPR1e
PpOPR1c
DsOPR1c
PpOPR1d
DsOPR1d
DsOPR1f
PpOPR1a
DsOPR1a
NeOPR1
AfOPR1
TaOPR2
ZmOPR4
TaOPR1
AtOPR1
AtOPR2
SlOPR1
PaOPR1
EhOPR1a
ScOPR1
SmOPR1b
SmOPR1c
SmOPR1a
EhOPR1b
DdOPR1b
LdOPR1a
PpOPR1b
DsOPR1e
PcOPR1
EhOPR1c
DdOPR1d
LdOPR1b
MpOPR1a
MpOPR1b
DdOPR1a
AaOPR1a
AaOPR1b
AaOPR1d
AaOPR1c
DsOPR1b
KnOPR1
Class I
0.5

**Fig. S6. Phylogenetic analysis of *OPR* genes**.

The rooted maximum-likelihood (ML) phylogenetic tree shown here was inferred from the amino acid sequences alignment of the full protein sequence from 18 plant species (Suppl. Table S1) by Dialing Multiple Alignment programme. The multiple alignment outcome was then processed with PhyML and TreDyn to obtain the phylogenetic tree. The bootstrap values from 1000 resamplings were calculated and the branch lengths are drawn to scale. Branches of the OPR sequences belonging to different clades are labelled in specific colours: spermatophytes (blue), monilophytes (green), lycophytes (red), bryophytes (orange) and algae (grey). The plant species analysed here are reported in Suppl. Table S1. OPR proteins containing a canonical peroxisomal signal are highlighted with a solid green triangle.

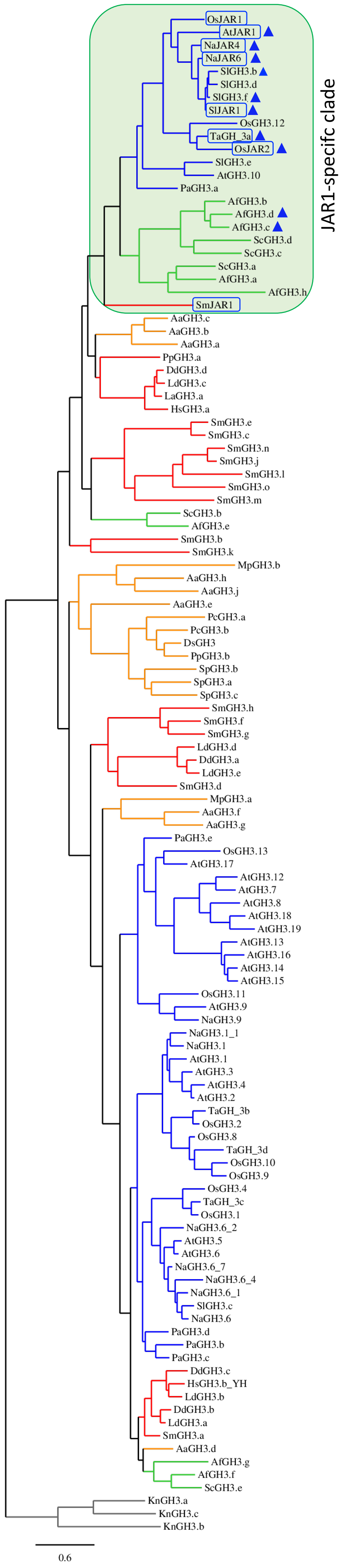

JAR1-specifc clade
OsJAR1
AtJAR1
NaJAR4
NaJAR6
SlGH3.b
SlGH3.d
SlGH3.f
SlJAR1
OsGH3.12
TaGH_3a
OsJAR2
SlGH3.e
AtGH3.10
PaGH3.a
AfGH3.b
AfGH3.d
AfGH3.c
ScGH3.d
ScGH3.c
ScGH3.a
AfGH3.a
AfGH3.h
SmJAR1
AaGH3.c
AaGH3.b
AaGH3.a
PpGH3.a
DdGH3.d
LdGH3.c
LaGH3.a
HsGH3.a
SmGH3.e
SmGH3.c
SmGH3.n
SmGH3.j
SmGH3.l
SmGH3.o
SmGH3.m
ScGH3.b
AfGH3.e
SmGH3.b
SmGH3.k
MpGH3.b
AaGH3.h
AaGH3.j
AaGH3.e
PcGH3.a
PcGH3.b
DsGH3
PpGH3.b
SpGH3.b
SpGH3.a
SpGH3.c
SmGH3.h
SmGH3.f
SmGH3.g
LdGH3.d
DdGH3.a
LdGH3.e
SmGH3.d
MpGH3.a
AaGH3.f
AaGH3.g
PaGH3.e
OsGH3.13
AtGH3.17
AtGH3.12
AtGH3.7
AtGH3.8
AtGH3.18
AtGH3.19
AtGH3.13
AtGH3.16
AtGH3.14
AtGH3.15
OsGH3.11
AtGH3.9
NaGH3.9
NaGH3.1_1
NaGH3.1
AtGH3.1
AtGH3.3
AtGH3.4
AtGH3.2
TaGH_3b
OsGH3.2
OsGH3.8
TaGH_3d
OsGH3.10
OsGH3.9
OsGH3.4
TaGH_3c
OsGH3.1
NaGH3.6_2
AtGH3.5
AtGH3.6
NaGH3.6_7
NaGH3.6_4
NaGH3.6_1
SlGH3.c
NaGH3.6
PaGH3.d
PaGH3.b
PaGH3.c
DdGH3.c
HsGH3.b_YH
LdGH3.b
DdGH3.b
LdGH3.a
SmGH3.a
AaGH3.d
AfGH3.g
AfGH3.f
ScGH3.e
KnGH3.a
KnGH3.c
KnGH3.b
0.6

**Fig. S7. Phylogenetic analysis of GH3 proteins.**

The phylogenetic analysis of the full protein sequence from 17 plant species (Suppl. Table S2) were aligned by MUSCLE Multiple Alignment programme to infer the unrooted maximum-likelihood (ML) tree. The multiple alignment outcome was then processed with PhyML and TreDyn to obtain the phylogenetic tree. The bootstrap values from 1000 resamplings were calculated and the branch lengths are drawn to scale. Branches of the GH3 sequences belonging to different clades are labelled in specific colours: spermatophytes (blue), monilophytes (green), lycophytes (red), bryophytes (orange) and algae (grey). The plant species analysed here are reported in Suppl. Table S2. GH3 proteins with experimentally proven *in vitro* or *in vivo* activity are highlighted in blue boxes. Proteins that conserved at least 3 of the 4 residues directly interacting with JA-Ile in the AtJAR1 structure are labelled with blue triangles. The GH3 proteins that closely cluster with AtJAR1 are highlighted in a green box.

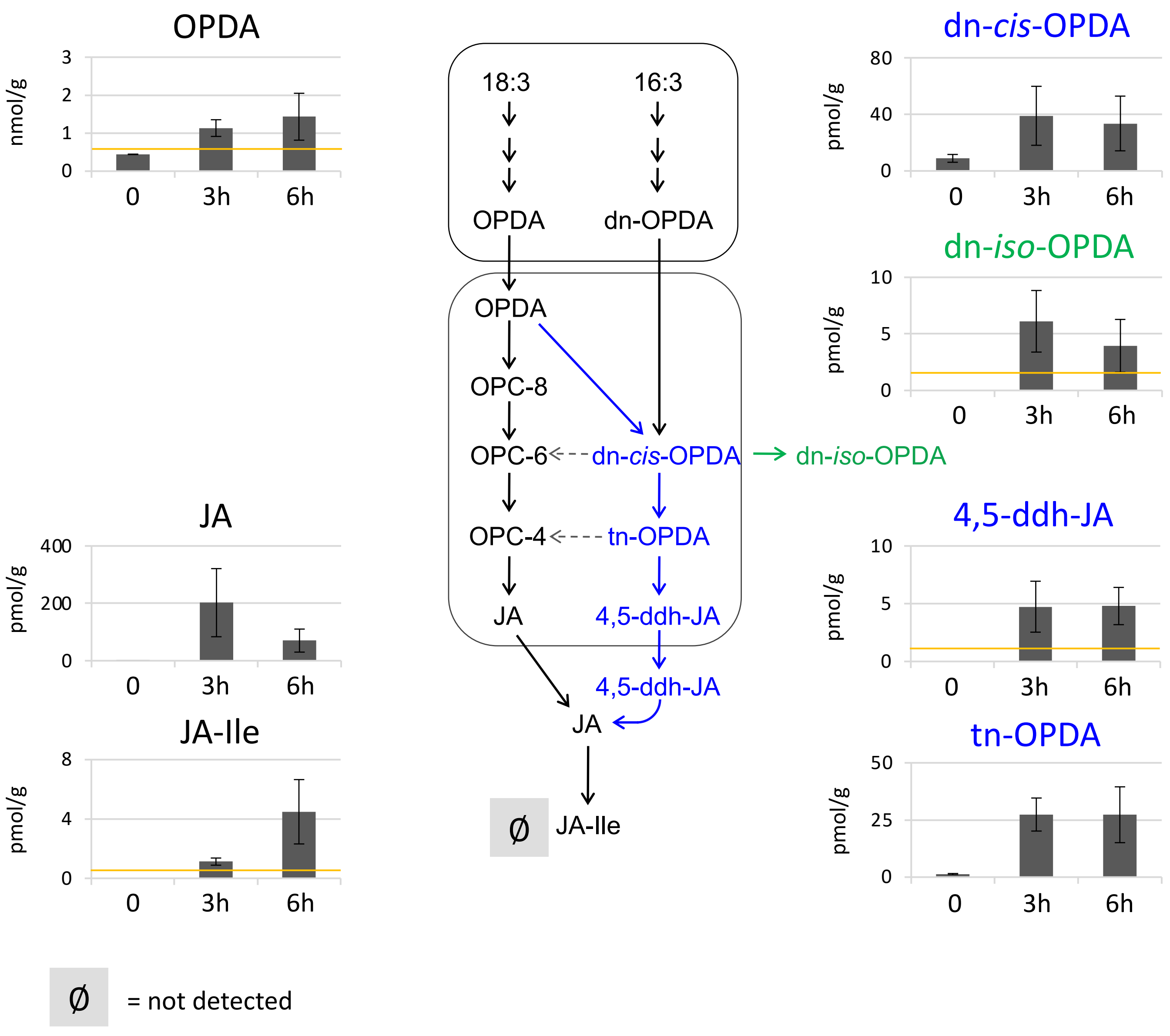


**Fig. S8. Accumulation of jasmonates in *Selaginella moellendorffii*.**

Time-course analysis of jasmonates accumulation after wounding in the lycophytes *Selaginella moellendorffii*. Wounded material was collected 3 and 6 hours after wounding. Unwounded plants (0) were included as control. Data shown as mean ± s.d. of three to four biological replicates. Experiments were repeated twice with similar results. An orange line shows the detection limit for OPDA (0.5 pmol/g), dn-*iso*-OPDA (1 pmol/g), 4,5-ddh-JA (1 pmol/g) and JA-Ile (0.5 pmol/g). In the center, a schematic representation of the jasmonate biosynthetic pathways is included. Black arrows define the canonical octadecanoid pathway, whereas blue arrows and molecules indicate the OPR3-independent pathway. Ø indicates compounds that were not detected.

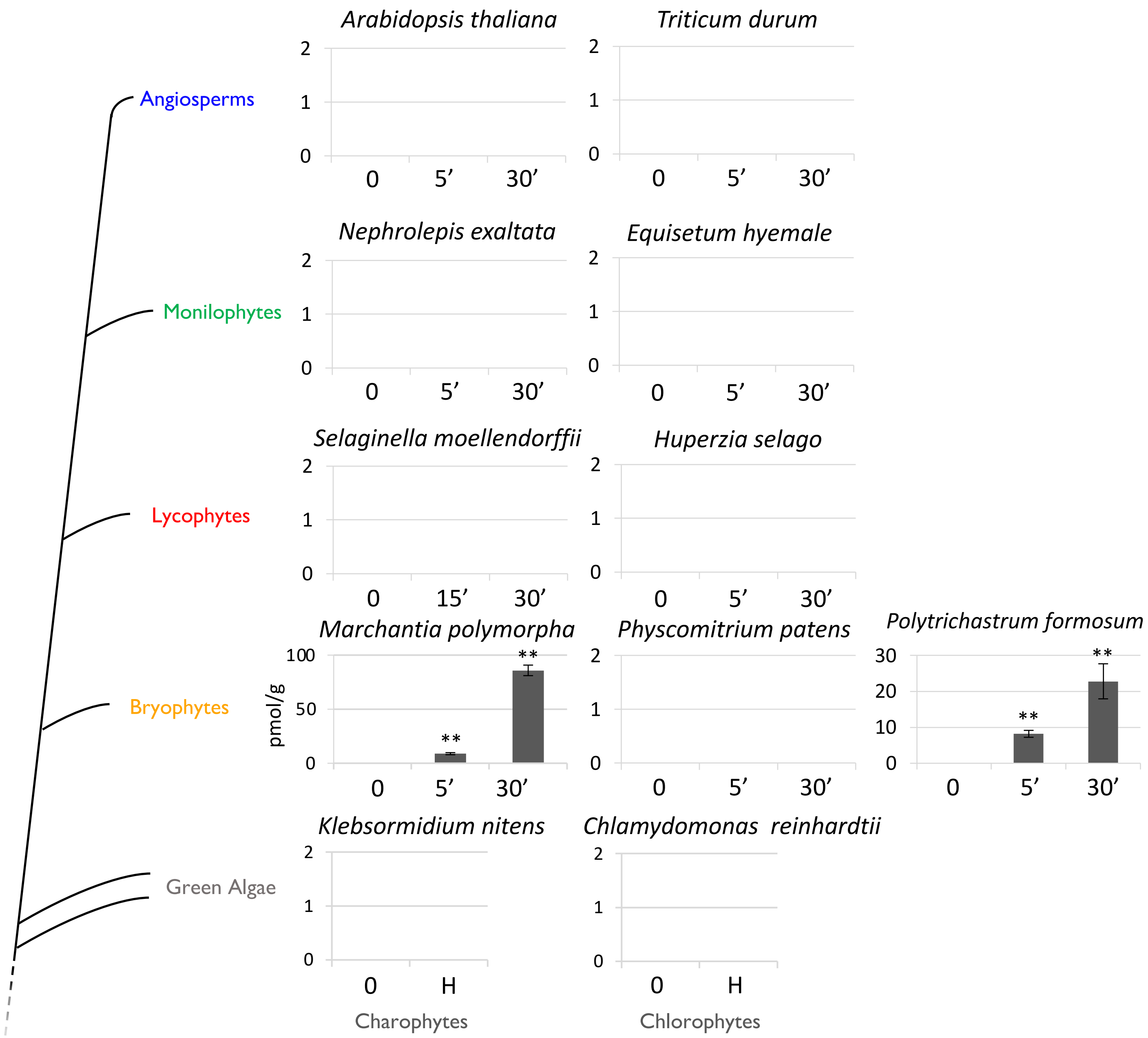


**Fig. S9. Accumulation of Δ[4]-dn-OPDA in representative plant species after wounding**.

Time-course accumulation of Δ[4]-dn-OPDA [pmol/fresh weight (g)] after wounding in several plant species. Data shown as mean ± s.d. of four biological replicates. Experiments were repeated 2 times with similar results. Asterisks indicate significant differences between wounded and unwounded samples according to a Student's t-test analysis (** p-val < 0.01). On the left, a schematic representation of a plant phylogenetic tree is included.

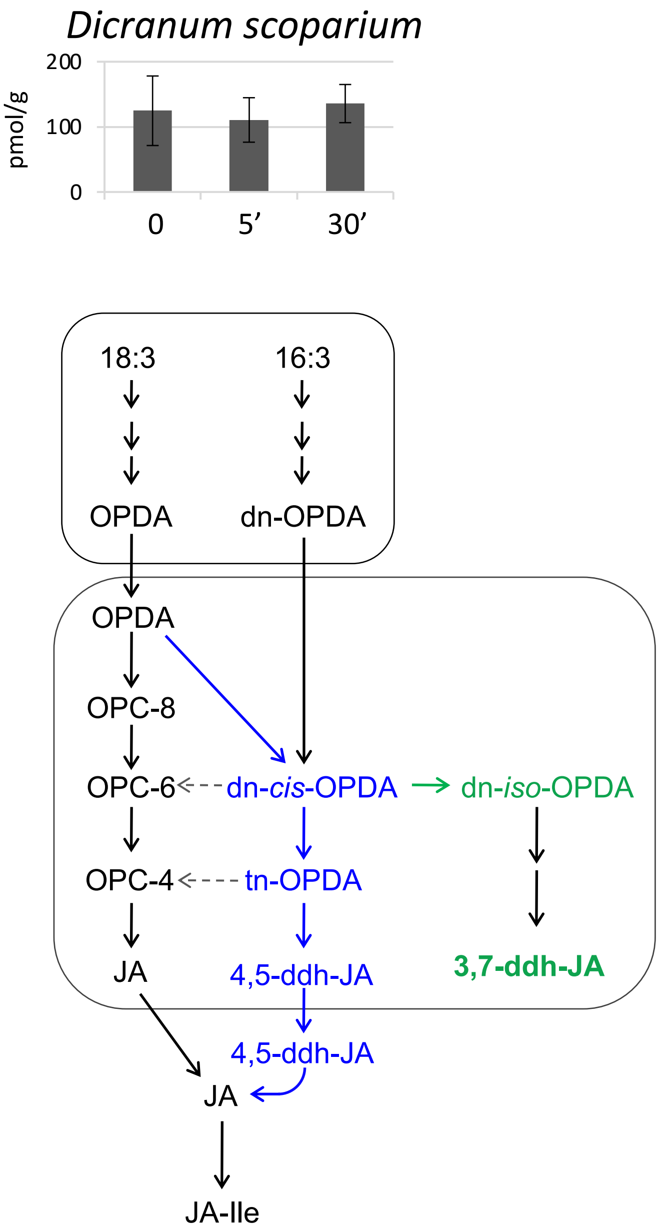


**Fig. S10. Accumulation of 3,7-ddh-JA in the moss *Dicranum scoparium*.**

Time-course analysis of 3,7-ddh-JA accumulation after wounding in the moss *Dicranum scoparium*. Damaged material was collected after the indicated times. Unwounded plants (0) were included as control. A schematic representation of the biosynthetic pathway of JA-Ile is reported on the bottom. Black arrows define the canonical octadecanoid pathway and blue arrows and molecules indicate the OPR3-independent pathway.

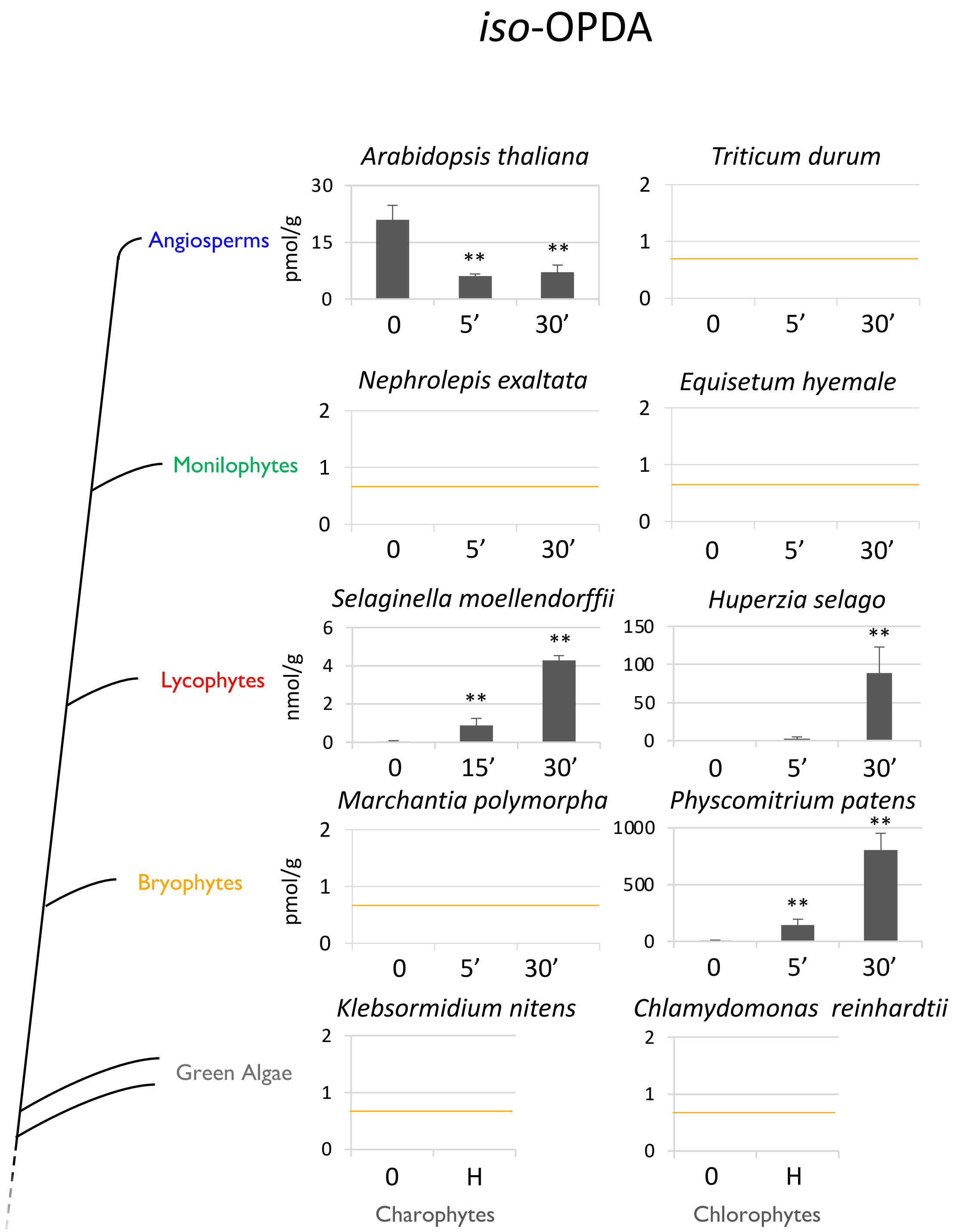


**Fig. S11. Accumulation of *iso*-OPDA in representative plant species after wounding**.

Time-course accumulation of *iso*-OPDA [pmol or nmol/fresh weight (g)] after wounding in several plant species. Data shown as mean ± s.d. of three or four biological replicates. Experiments were repeated 2 to 4 times with similar results. Asterisks indicate significant differences between wounded and unwounded samples according to a Student's t-test analysis (** p-val < 0.01). An orange line shows the detection limit (0.7 pmol) in the case of plants in which *iso*-OPDA was not detected. On the left, a schematic representation of a plant phylogenetic tree is included.
